# EpiMix: an integrative tool for epigenomic subtyping using DNA methylation

**DOI:** 10.1101/2023.01.03.522660

**Authors:** Yuanning Zheng, John Jun, Kevin Brennan, Olivier Gevaert

## Abstract

DNA methylation (DNAme) is a major epigenetic factor influencing gene expression with alterations leading to cancer, immunological, and cardiovascular diseases. Recent technological advances enable genome-wide quantification of DNAme in large human cohorts. So far, existing methods have not been evaluated to identify differential DNAme present in large and heterogeneous patient cohorts. We developed an end-to-end analytical framework named “EpiMix” for population-level analysis of DNAme and gene expression. Compared to existing methods, EpiMix showed higher sensitivity in detecting abnormal DNAme that was present in only small patient subsets. We extended the model-based analyses of EpiMix to cis-regulatory elements within protein-coding genes, distal enhancers, and genes encoding microRNAs and lncRNAs. Using cell-type specific data from two separate studies, we discovered novel epigenetic mechanisms underlying childhood food allergy and survival-associated, methylation-driven non-coding RNAs in non-small cell lung cancer.

## Main text

DNA methylation (DNAme) is one of the major epigenetic marks in humans. It is defined as the addition of a methyl (CH_3_) group to DNA that occurs primarily at the cytosine of cytosine-guanine dinucleotide (CpG) sequence. DNAme regulates various biological processes by affecting gene expression, and aberrant DNAme plays a critical role in the development and progression of many human diseases^1–3^. Recent experimental methods based on microarrays or next-generation sequencing have enabled genome-wide quantification of DNAme at single-nucleotide resolution. Due to its quantitative and cost-effective nature, microarray-based technology has emerged as the method of choice for profiling DNAme in large human cohorts. For example, The Cancer Genome Atlas (TCGA) project has used the microarray technology to generate DNAme profiles in over 10,000 specimens representing 33 cancer types. The Gene Expression Omnibus database (GEO) and other public repositories also host a large number of DNAme datasets across cancers and other complex diseases.

Over the last decade, a number of computational approaches have been developed to identify genes that are abnormally methylated in human diseases. Some methods are tailored to the analysis of DNAme data from bisulfite sequencing^4–7^, while others are designed for array-based data or can be adapted to both data platforms^8–12^. Many existing methods identify differentially methylated loci by comparing all samples from an experimental group versus samples in a control group. This type of comparison works well when the experimental population is assumed to be homogenous. However, when the study population is large, abnormal DNAme may be present in only a subset of the patients, and this intra-population variation has been observed in cancers and many other diseases^13–15^. In cases where abnormal DNAme occurred in only a small subset of the patients, existing methods are not capable of capturing the signals of differential methylation. Therefore, there is a critical need to use a statistical approach to model the distribution of DNAme in large patient cohorts, and to identify the patient subsets with differential DNAme profiles. This epigenetic subtyping can be essential to improve personalized diagnosis, treatment and drug discovery.

Furthermore, gene expression in mammalian cells is a result of a complex process coordinated by a broad range of genomic regulatory elements^16,17^. In many studies, CpG sites were mapped to genes based on linear genomic proximity. This mapping logic assumes that the transcriptional activity can be affected only when the genes are overlapped or close to the differentially methylated sites. However, emerging evidence has shown that distal enhancers, which may locate at a great linear genomic distance from their target genes, play a critical role in orchestrating spatiotemporal gene expression programs^18^. Abnormal DNAme at enhancers was frequently reported in cancers and many other diseases^19,20^. Therefore, the analysis of enhancer methylation can improve our understanding of how gene expression is regulated across physiological and pathological conditions.

Existing computational tools focus on the DNAme analysis of protein-coding genes. Besides protein-coding genes, non-coding RNAs, such as microRNAs (miRNAs) and long non-coding RNAs (lncRNAs), play an important role in regulating cell functions^21,22^. Recent studies have shown that DNAme is a major epigenetic mechanism regulating non-coding RNA expression^23,24^. With existing methods, it is challenging to decipher how DNAme regulates non-coding RNA expression.

Here, we present EpiMix, a comprehensive analytical framework for population-level analysis of DNAme and gene expression. EpiMix utilizes a model-based computational approach to identify abnormal DNAme at diverse genomic elements, including cis-regulatory elements within or surrounding protein-coding genes, distal enhancers, and genes encoding miRNAs and lncRNAs. In two separate studies, we showed that EpiMix identified novel methylation-driven pathways in T cells from childhood food allergy and methylation-driven non-coding RNAs in non-small cell lung cancer patients. To improve usability, we disseminated EpiMix’s algorithms in Bioconductor^25^, enabling end-to-end DNAme analysis. Furthermore, we developed a web tool for interactive exploration and visualization of EpiMix’s results (https://epimix.stanford.edu). Overall, EpiMix can be used to discover novel epigenetic biomarkers for disease subtypes and therapeutic targets for personalized medicine.

## Results

### Overview of EpiMix Workflow

EpiMix is an end-to-end analytical framework for modeling DNAme at diverse genomic elements and for identifications of differential DNAme associated with gene expression. The EpiMix framework consisted of four functional modules: (1) data downloading, (2) preprocessing, (3) DNAme modeling and (4) functional analysis (**Fig.1**). To analyze DNAme at functionally diverse genomic elements, we implemented four alternative analytic modes: “Regular,” “Enhancer”, “miRNA” and “lncRNA.” Both the Regular and Enhancer modes aimed to detect differential DNAme associated with the expression of protein-coding genes. The Regular mode analyzed DNAme sites within or immediately surrounding the genes, while the Enhancer mode specifically analyzed DNAme at distal enhancers. The miRNA and lncRNA modes were built for the detection of DNAme affecting the expression of miRNAs and lncRNAs. After the methylation-driven genes were identified, users could perform comprehensive exploratory analyses using the functional analysis module. The functional analysis module was built with both in-house developed methods and integrating existing computational tools to enable diverse functional analyses and visualization of the differential DNAme.

**Fig.1.**
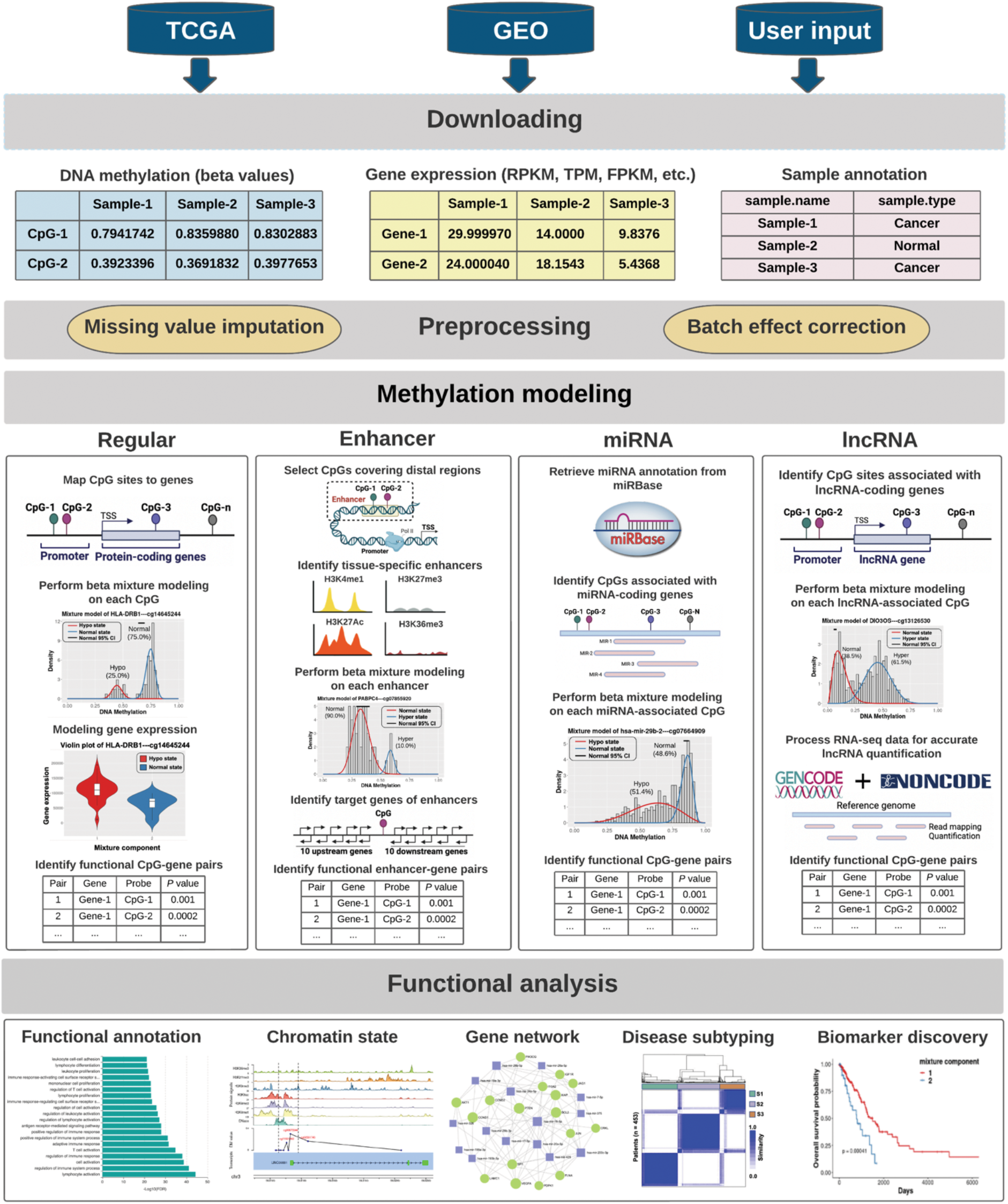
Overview of EpiMix workflow. EpiMix includes four modules: Downloading, Preprocessing, Methylation modeling and Functional analysis. Data from public repositories (i.e., TCGA and GEO) can be automatically downloaded and preprocessed by EpiMix. Alternatively, users can input their own custom datasets. The preprocessing module includes functions for quality control, batch effect normalization, and missing value imputation. To model DNAme, EpiMix enables four alternative analytic modes: Regular, Enhancer, miRNA and lncRNA. Each mode uses a custom algorithm to analyze DNAme at a specific type of genomic element. One major output from the methylation modeling is a matrix of functional CpG-gene pairs, illustrating the differentially methylated CpGs whose DNAme states were associated with gene expression. After the differentially methylated genes have been identified, users can perform diverse analytical tasks with EpiMix’s functional analysis module. This includes pathway enrichment analysis, genome-browser style visualization, gene regulatory network analysis, epigenetic biomarker discovery and identification of methylation-associated disease subtypes.

### Identifications of abnormal DNAme present in small sample subsets

To assess the sensitivity of EpiMix in identifications of differential DNAme that was present in only specific patient subsets, we performed simulation experiments. We used a dataset that jointly profiled DNAme data and messenger RNA abundance in human naïve CD4+ T cells^26^. The dataset contains quiescent T cells and antigen-activated T cells from 103 human subjects. The DNAme data were obtained from Infinium MethylationEPIC array, and the messenger RNA expression data were obtained from RNA-Seq. We randomly sampled a subset of CpGs (n = 300) from the quiescent group as baselines, such that the average beta values of the selected CpGs ranged from 0.1 to 0.9. Then, for each CpG, we randomly selected a subset of samples from the activation group and combined them with the baseline group (**Fig.2a** and **Methods**), such that the final proportions of samples from the activation group in the combined dataset ranged from 3% to 50%, and the mean differences in beta values between the activated and the baseline samples ranged from 0.1 to 0.7. We then compared the DNAme of the synthetic populations to the baseline population (**Fig.2a**).

**Fig.2.**
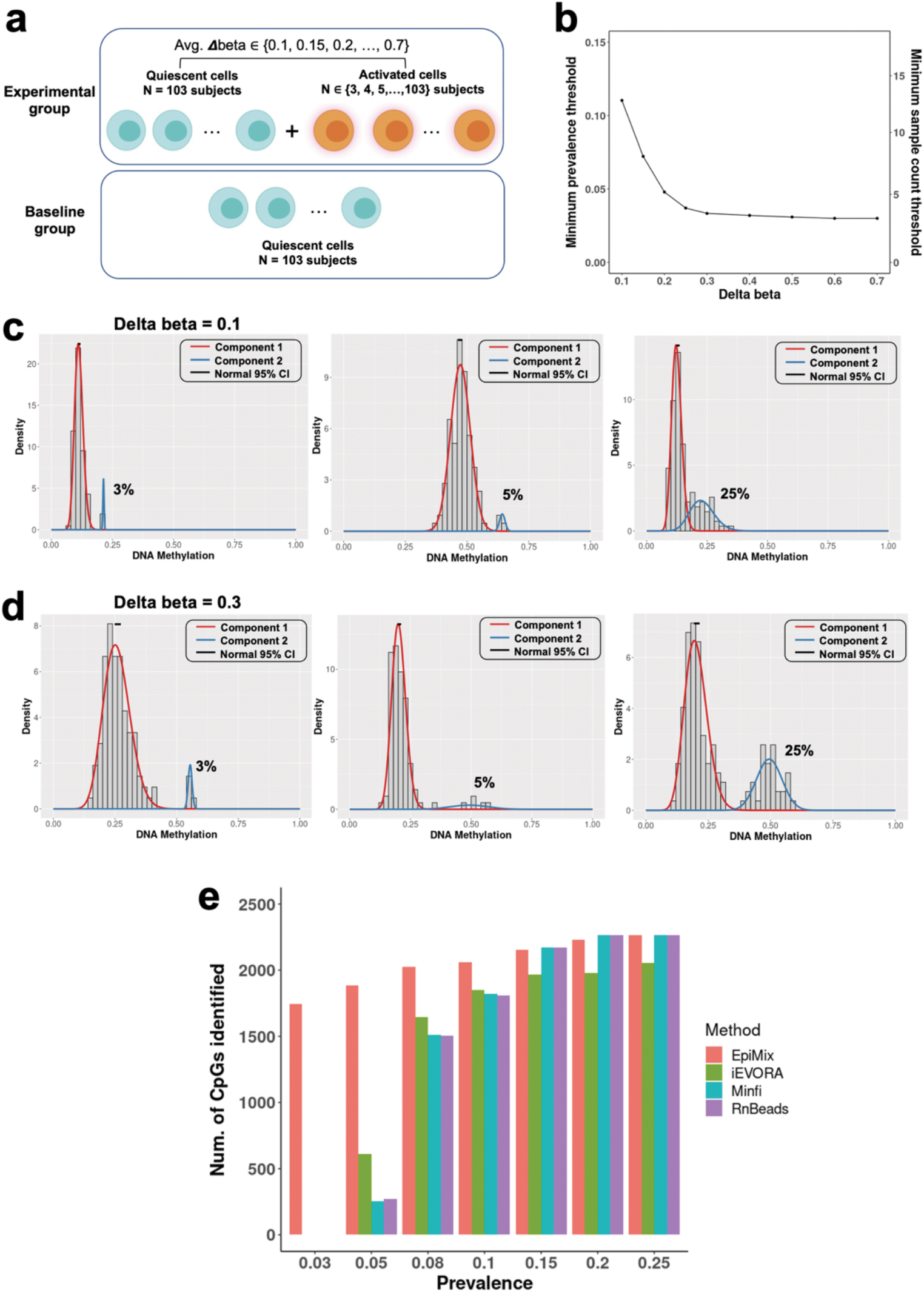
**a**, Design of the simulation study. The dataset contained experimentally purified naïve CD4+ T cells from 103 human subjects. Cells from each subject were divided into half and either activated with the T-cell antigen or left resting in the media. The baseline group contained quiescent samples from all 103 subjects. The experimental group contained quiescent samples from all subjects and the antigen-activated samples from *N* subjects, where *N* ranged from 3 to 103. We compared the DNAme of the experimental group to the baseline group and tested whether EpiMix can detect the signals of differential methylation. **b**, Correlation between the delta beta values and the minimum detection threshold for the prevalence (left axis) and actual count (right axis) of the activated samples in the experimental group. The simulation was repeated 300 times using a different CpG site at each time, and the mean detection threshold was shown. **c**, Density plots showing the mixture models when delta beta was 0.1 and the differential methylation was present in 3%, 5% and 25% of the experimental group. **d**, Density plots showing the mixture models when delta beta was 0.3 and the differential methylation was present in 3%, 5%, and 25% of the experimental group. **e)** Number of differentially methylated CpGs detected by different methods when the differential methylation was present in from 3% to 25% of the population. For all methods, the same set of CpGs were used, and the total number of CpGs at each prevalence was 2,700.

We found that the sensitivity of EpiMix was determined by the magnitude of differences in DNAme between the quiescent and the activated subjects. When the delta beta was 0.1, EpiMix detected differential DNAme that was present in 3% to 25% of the synthetic population, with a mean minimum detection threshold of 11.0% (absolute sample count = 13) (**Fig.2b, c**). When the delta beta was 0.2 or higher, the minimum detection threshold ranged from 3% to 10%, with a mean threshold of 3.4% (absolute sample count = 4) (**Fig.2b, d**). These results indicated that EpiMix was able to detect abnormal DNAme that was present in only small subsets of a tested population, and the sensitivity was positively correlated with the magnitude of differences in DNAme.

Next, we compared the performance of EpiMix with other existing methods in identifications of differential DNAme, including Minfi^10^, iEVORA^27^ and RnBeads^12,28^. When the differential DNAme was present in 3% of the population, EpiMix detected the differential methylation signals at 1,747 CpG sites, whereas the other methods did not capture any differential DNAme (**Fig.2e)**. When the differential DNAme was present in 5% of the population, EpiMix identified 3.1 times more differentially methylated CpGs than iEVORA, and 3.6 times more CpGs than Minfi and RnBeads. Minfi and RnBeads only detected CpGs with high magnitude differences in DNAme, with an average delta beta of 0.6. In contrast, EpiMix detected CpGs with delta beta ranging from 0.1 to 0.7, with an average threshold of 0.3. When the prevalence of differential DNAme was 15% or higher, EpiMix detected similar numbers of CpGs to the other three methods. These results indicated that EpiMix had higher sensitivity to detect differential DNAme that was present in only small sample subsets.

### Modeling of DNA methylation at *cis*-regulatory elements within protein-coding genes

To test the Regular mode of EpiMix, we used the complete, real dataset from antigen-activated T cells and quiescent T cells (n = 103 subjects per group)^26^. In the activated T cells, 1,090 CpGs were differentially methylated compared to the quiescent cells. Integrative analysis with RNA-seq data showed that the differentially methylated CpGs were functionally associated with the expression of 748 protein-coding genes (**Supplementary Table 1**). Of the differentially methylated CpGs, 746 (68.4%) CpGs associated with 504 genes were hypomethylated and 327 (30.0%) CpGs associated with 238 genes were hypermethylated (**Fig.3a**). This result indicated that antigens induced a widespread loss of DNAme. Gene ontology (GO) analysis showed that the hypomethylated genes were associated with lymphocyte proliferation (e.g., *CCND2, CCND3, CDK6, CDK14*), T cell activation (e.g., *BCL2, CCL5, HLA-DPA1, HLA-DRB1*), glycoprotein biosynthesis (e.g., *AGO2, ALG9, B3GNT5, B4GALT5*) and cytokine receptor activity (*IL1R1, IL1R2, IL21R, IL23R*) (**Supplementary Table 2**). This result confirmed that EpiMix identified differential DNAme associated with T cell activation.

**Fig.3.**
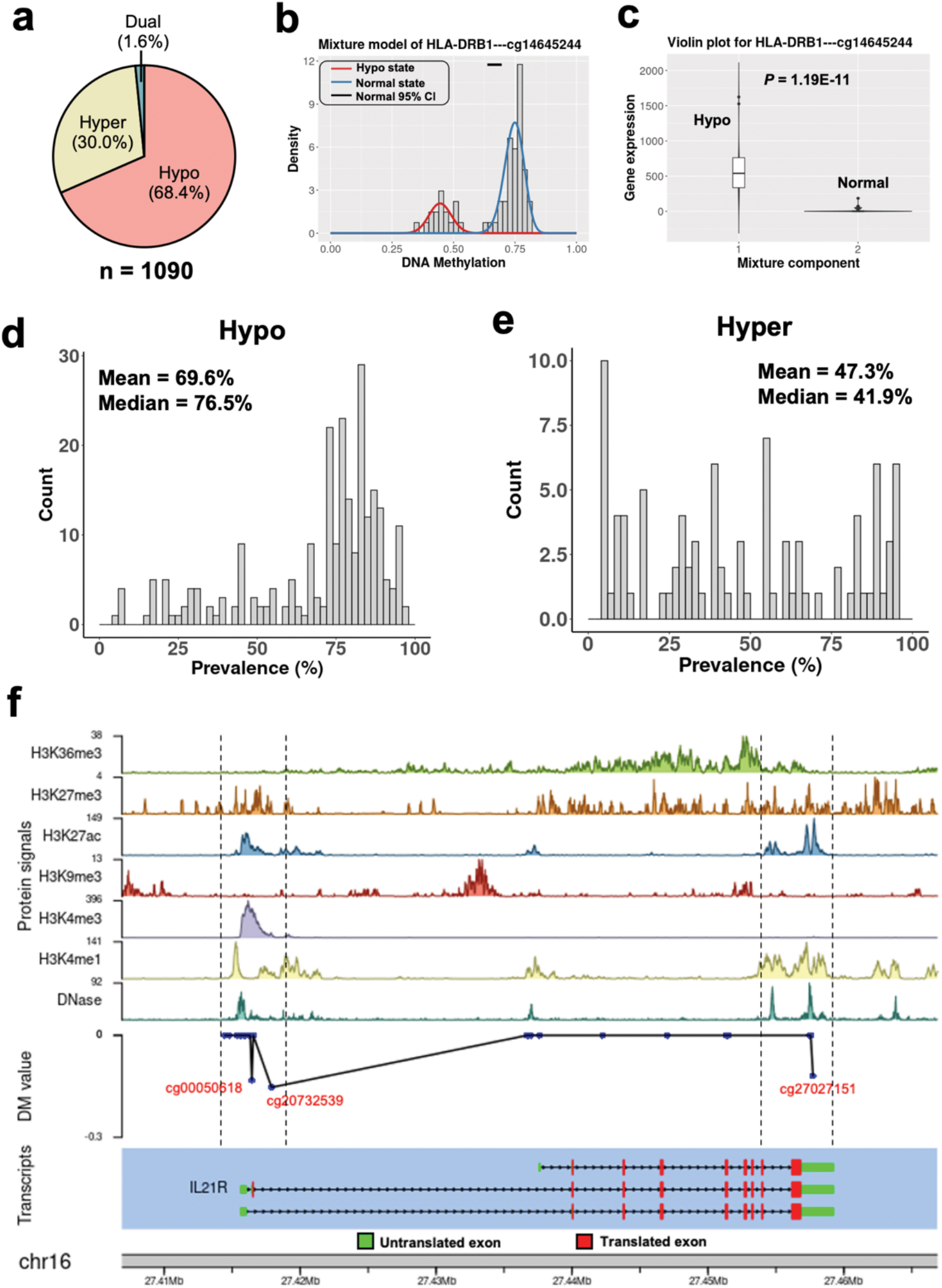
Identifications of differential DNAme resulting from antigen-induced T cell activation. **a**, Proportions of the hypo-, hyper- and dual methylated CpGs in antigen-activated T cells. The dual methylated CpGs refer to the CpGs that were hypomethylated in some individuals, while hypermethylated in some other individuals. **b**, Mixture model of a CpG associated with the *HLA-DRB1* gene, and **c**, *HLA-DRB1* gene expression levels in different mixtures. Red indicates hypomethylation (n = 26), while blue indicates normal methylation (n = 77). Gene expression levels were compared with Wilcoxon rank-sum test. **d-e**, Density plots showing the prevalence distribution of the d) hypo-and e) hyper-methylated CpGs **f**, Genome-browser style visualization of the chromatin state, DM values, and transcript structure of the *IL21R* gene. The hypomethylated CpGs were labeled in red. The differential methylation (DM) value represents the mean difference in beta values between the hypomethylated subjects versus the normally methylated subjects. DM = 0: normal methylation; DM < 0: hypomethylation.

Many of the CpGs were differentially methylated in only a subset of the patients. For instance, the *Human Leukocyte Antigen DRB1* (*HLA-DRB1*) gene was hypomethylated in the antigen-activated T cells from 25% of the subjects, whereas the majority (75%) of the subjects had a normal methylation state similar to the quiescent T cells (**Fig.3b**). As expected, gene expression levels of *HLA-DRB1* were significantly increased in the hypomethylated compared to the normally methylated subjects (**Fig.3c**). Overall, the prevalence of hypomethylation ranged from 5.9% - 100%, with a mean prevalence of 69.6% (**Fig.3d**). The prevalence of hypermethylation ranged from 5.8% - 100%, with a mean prevalence of 47.3% (**Fig.3e**). These results indicated that the antigen-induced response in T cells varied between different individuals.

We next investigated the genomic distribution of the differentially methylated CpGs. Thirty-nine percent (39.5%) of the CpGs were located at the promoters, and 56.4% were located at introns (**Supplementary Fig.1a**). Using publicly available chromatin immunoprecipitation-sequencing (ChIP-seq) data of human naïve CD4+ T cells, we found that the abnormal DNAme was significantly enriched at active promoters marked by H3K4me3 and H3K27ac, active enhancers marked by H3K4me1, and to a lesser extent, actively transcribed gene bodies marked by H3K36me3 (**Supplementary Fig.1b**). These results demonstrated that EpiMix was able to identify aberrant DNAme at lineage-defining *cis*-regulatory elements.

To allow users to investigate the genomic locations and chromatin states associated with the differentially methylated sites, EpiMix enables genome browser-style visualization. We illustrated this functionality with hypomethylation in two regions of the interleukin-receptor gene *IL21R* (**Fig.3f)**. The first region was located at the promoter, which overlapped with DNase I hypersensitivity sites and activating histone modifications (i.e., H3K4me1, H3K4me3 and H3K27ac). The second region was located at the three-prime untranslated region, enriched with histone modifications marking for active enhancers (i.e., H3K4me1 and H3K27ac). In concordance with this DNA hypomethylation, *IL21R* expression levels were significantly increased (**Supplementary Table 1**, Wilcoxon rank-sum test, *P* < 3.19E-08).

#### Identification of functional DNA methylation at distal enhancers in food allergy

To demonstrate the Enhancer mode of EpiMix, we used the same CD4+ T cell dataset^26^. In this dataset, 82 human subjects were diagnosed with food allergy and 21 subjects were non-allergic controls. The differential response of T cells to antigen-induced activation between different individuals may be associated with the allergic status. We then characterized allergy-associated changes in DNAme by comparing antigen-activated T cells from the allergic patients to those from the non-allergic controls. Using a permutation approach (**Supplementary Fig.2** and **Methods**), we identified 107 differentially methylated enhancers that were functionally linked to the expression of 119 genes. The number of target genes of each enhancer ranged from 1 to 3, resulting in 131 significant enhancer-gene pairs (**Supplementary Table 3**). This result is consistent with the previous studies showing that enhancers typically loop to and are associated with the activation of 1 to 3 promoters^29,30^. Of the functional enhancers, 21/107 (19.6%) enhancers associated with 24 genes were hypomethylated, 82/107 (76.7%) enhancers associated with 92 genes were hypermethylated (**Fig.4a**). This result indicated that there was a global gain of DNAme at enhancers in food allergy.

**Fig. 4.**
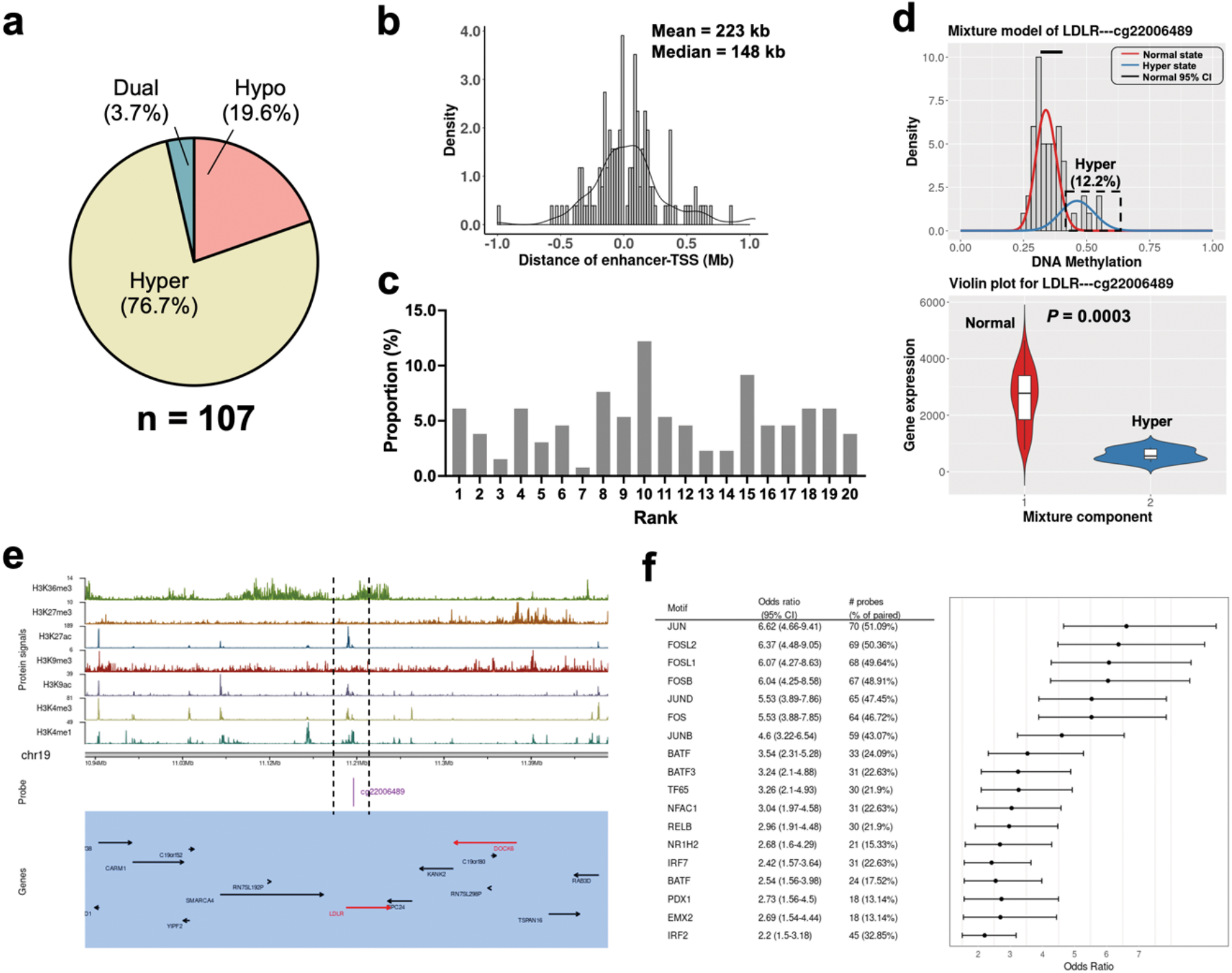
Identifications of differentially methylated enhancers associated with food allergy. **a**, Proportions of the hypo-, hyper- and dual methylated enhancers in children with food allergy. **b**, Distribution of the linear genomic distance between enhancers and their gene targets. **c**, For each functional enhancer, the 20 adjacent genes were ranked by genomic distance. Bars show the proportions of the functionally linked genes in each rank. **d**, Mixture model of the *LDLR* gene (top panel) and *LDLR* gene expression levels in different mixtures (bottom panel). Red indicates normal methylation (n = 72), while blue indicates hypermethylation (n = 10). Gene expression levels were compared by Wilcoxon rank-sum test. **e**, Integrative visualization of the chromatin states and the adjacent genes of the hypermethylated enhancer shown in panel d. The genes in the functional CpG-gene pairs are shown in red, while the others are shown in black. **f**, Enriched TF motifs and odds ratios for the differentially methylated enhancers. To find significantly enriched motifs, we used all the distal CpGs as the background and the functional enhancers as the targets.

The genomic distance between enhancers and their target genes ranged from 4.5 kb to 1.7 Mb, with a median distance of 148 kb (**Fig.4b**). In a previous study, Jin et al. used high-throughput chromosome conformation capture (Hi-C) assay to investigate promoter-enhancer interactions and demonstrated that approximately 25% of the enhancer-promoter pairs are within a 50 kb range and approximately 57% spans 100 kb or greater genomic distance, with a median distance of 124 kb^31^. Another study by Rao et al. showed that the distance between enhancers and promoters spans from 40 kb to 3 MB, with a median distance of 185 kb^32^. Our data agree with these experimentally generated results. To further characterize the enhancer-gene linkage, we investigated how often did the functional enhancers associate with the nearest gene promoter. We ranked the 20 adjacent genes of each enhancer by their genomic distance to the enhancer. **Fig.4c** showed that only 6.1% of the times did the enhancer associate with the nearest promoter, whereas the majority of the enhancers skipped one or more intervening genes to associate with promoters farther away. In line with this result, a previous study using the chromosome 5C assay showed that only ∼7% of the time did the distal elements loop to the promoter of the nearest gene, whereas the majority of enhancers bypass the nearest promoter and loop to promoters farther away^33^. These results confirmed that EpiMix identified true distal *cis*-regulatory events.

The genes linked to the differentially methylated enhancers were related to the lipid metabolism (*LDLR, CAT, LPIN2, SREBF1, PIK3C2B*) and T cell activation (*CASP3, MALT, PRKCZ, SMAD3*). **Fig.4d** showed that the enhancer linked to the *LDLR* gene was hypermethylated in 12.2% of the allergic patients, and the gene expression of *LDLR* was significantly decreased in the hypermethylated patients. Integrative visualization (**Fig.4e**) showed that the hypermethylated enhancer overlapped with the Dnase I hypersensitivity site and was enriched with histone modifications marking for active enhancers, including H3K4me1 and H3K27ac, and to a lesser extent, H3K4me3 and H3K9ac. The *LDLR* gene encodes a low-density lipoprotein receptor that transports cholesterol from the blood into the cell, which plays a critical role in regulating T cell lipid metabolism^34^. Our results suggested that T cells from a small subset of the allergic patients may have an abnormal lipid metabolic profile due to enhancer hypermethylation.

Enhancers are enriched for sequences bound by site-specific transcription factors (TFs). Hypermethylation of enhancers suppresses gene transcription by decreasing the binding affinity of TFs^35,36^. We then carried out motif enrichment analysis of the differentially methylated enhancers. We identified significant enrichment of binding sites for Jun-related factors (JUN, JUND), Fos-related factors (FOS, FOSL1, FOSL2, FOSB), BATF-related factors (BATF, BATF3), and Interferon-regulatory factors (IRF2, IRF5, IRF7) (**Fig.4f** and **Supplementary Table 4**). These results agree with the evidence showing that Jun-related factors, BATF-related factors and Interferon-regulatory factors play a critical role in regulating the immune gene activation in T cells, and dysregulation of their activity causes aberrant immune response^37,38^. Our results demonstrated that the abnormal DNAme at enhancers affected the target gene response of these TFs and increased the subsequent risk for developing food allergy.

#### Identification of methylation-driven miRNAs in human lung cancer

Similar to protein-coding genes, miRNA-coding genes are transcriptionally regulated by DNAme^39,40^. To demonstrate the miRNA mode of EpiMix, we used a lung adenocarcinoma dataset containing DNAme and miRNA expression profiles of 457 tumors and 32 adjacent normal tissues^41^. The DNAme data were acquired from the HM450 array, and the gene expression data were obtained from high-throughput microRNA sequencing (miRNA-Seq).

Both tumors and normal tissues from the lung are composed of multiple cell types, majorly including epithelial cells, fibroblasts, hematopoietic cells and endothelial cells. Studies have shown that DNAme profiles are cell-type specific^42,43^. When using data collected at the tissue (“bulk”) level for DNAme analysis, the differential DNAme may result from variations in cell-type proportions between different individuals. To resolve the confounding effects from intra-tumoral heterogeneity, we used previously validated computational methods to decompose tissue compositions and to infer cell-type-specific methylomes and transcriptomes (**Supplementary Fig. 3** and **Methods**)^44,45^. We then applied EpiMix to the deconvoluted data of each individual cell type. In epithelial cells, we identified 272 differentially methylated CpGs functionally associated with the expression of 92 miRNA genes (**Fig.5a** and **Supplementary Table 5**). In fibroblasts, we found 12 hypomethylated CpGs functionally associated with the expression of 3 miRNA genes (**Supplementary Fig. 4a-b**). Although we discovered 9 differentially methylated CpGs in hematopoietic cells and 6 CpGs in endothelial cells, none of the differential DNAme were functionally correlated with gene expression. We further compared the differentially methylated gene lists identified using data from bulk tissues versus the ones using individual cell types. Over 80% of the differentially methylated genes identified in epithelial cells could also be identified using data from bulk tissues (**Supplementary Fig. 4a-b**). These results demonstrated that, although tumors are composed of multiple cell types, the majority of differential methylation events occurred in epithelial cells.

We next focused our analysis on the deconvoluted data of epithelial cells. Of the 272 differentially methylated CpGs, 138 (50.8%) CpGs associated with 66 genes were hypomethylated and 55 (20.2%) CpGs associated with 37 genes were hypermethylated. Sixty-five percent (63.6%) of the functional CpGs were located at the promoters, and this proportion was significantly higher than randomly selected CpGs (**Supplementary Fig.1c**, Fisher’s exact test, *P* = 0.003). Using publicly available ChIP-seq data of lung, we further determined that the differentially methylated regions were enriched with histone modifications (i.e., H3K27ac, H3K4me1 and H3K4me3) marking for actively transcribed promoters and enhancers (**Supplementary Fig.1d**). The prevalence of hypomethylation ranged from 1.1% to 66.7%, with a mean prevalence of 18.0% (**Fig. 5b**). Similarly, the prevalence of hypermethylation ranged from 2.6% to 83.7%, with a mean prevalence of 24.9% (**Fig. 5c**). These results indicated that the majority of differential DNAme associated with miRNA genes occurred in less than 25% of the patient population.

**Fig. 5.**
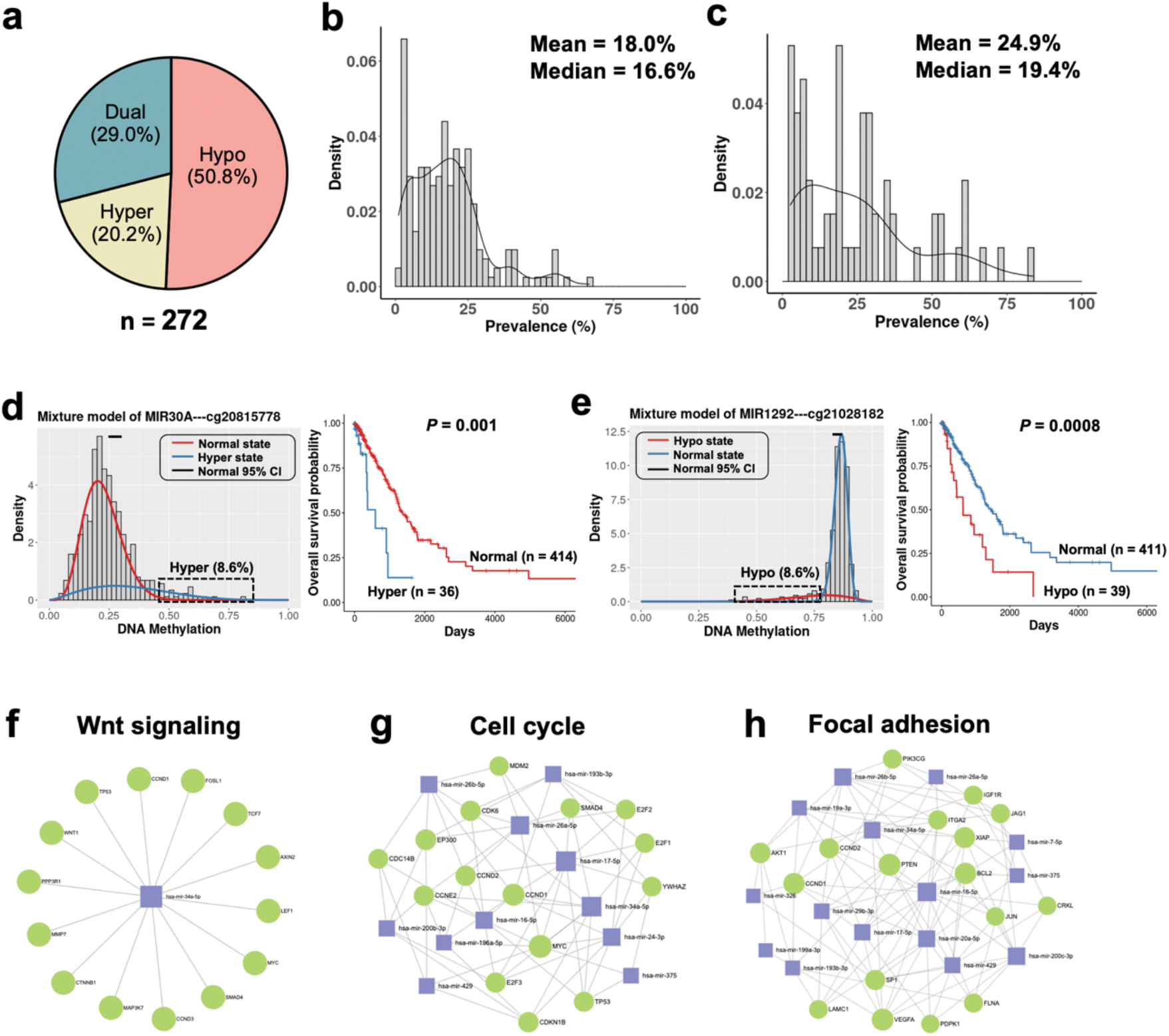
Identifications of differentially methylated miRNA-coding genes in human lung cancers. **a**, Proportions of the hypo-, hyper- and dual methylated CpGs of miRNAs in lung cancer. **b-c**, Density plots showing the prevalence distribution of the differentially methylated miRNAs in lung cancers (n = 457), **(b)** prevalence of hypomethylation and **(c)** prevalence of hypermethylation. **d**, Mixture model of the *MIR30A* gene (left panel) and Kaplan-Meier survival curves of patients in different mixtures (right panel). Red indicates normal methylation and blue indicates hypermethylation. Gene expression levels were compared by Wilcoxon rank-sum test. **e**, Mixture model of the *MIR1292* gene (left panel) and Kaplan-Meier survival curves of patients in different mixtures (right panel). Red indicates hypomethylation and blue indicates normal methylation. **f-g-h**, Network visualization of (**f**) the gene targets of miR-34a, (**g**) differentially methylated miRNAs related to the cell cycle pathway, and (**h**) focal adhesion pathway. Blue squares: miRNAs, green circles: protein-coding genes targeted by miRNAs.

MicroRNAs play an important role in regulating cell proliferation, invasion and cancer metastasis^46,47^. We next investigated whether the DNAme of miRNAs were associated with patient survival. Of the 92 methylation-driven miRNAs, we identified 22 miRNAs whose methylation states were significantly correlated with patient survival (**Supplementary table 6**, log-rank test, *P* < 0.05). Half (11/22, 50%) of the survival-associated miRNAs were hypomethylated and the others (11/22, 50%) were hypermethylated. Some of the miRNAs, such as *MIR29C*^48^, *MIR30A*^49^, *MIR34A*^50^ and *MIR148A*^51^, were known to be associated with lung cancer survival. For instance, MIR30A, a tumor suppressor miRNA^49^, was hypermethylated in 8.6% of the patients, and the hypermethylated patients showed a significantly worse survival than the normally methylated patients (**Fig.5d**, Hazard Ratio = 1.50, *P* = 0.001). In addition, EpiMix identified many new survival-associated miRNAs. For instance, *MIR1292* was hypomethylated in 8.6% of the patients, and the hypomethylated patients showed significantly worse survival (**Fig.5e**, Hazard Ratio = 1.39, *P* = 0.0008). These results demonstrated that EpiMix was able to identify survival-associated miRNAs that were differentially methylated in only small subsets of the patients, and this feature can be used to discover novel epigenetic biomarkers for prognosis.

To gain systematic insight into the biological functions of the methylation-driven miRNAs, we queried miRTarBase^52^ to obtain experimental validated target genes of the miRNAs. We then performed pathway analyses of the target gene list. The differentially methylated miRNAs were related to Wnt signaling pathway, cell cycle, p53 signaling, focal adhesion and apoptosis (**Fig.5f-h** and **Supplementary Table 7**). These results provided mechanistic insights into how abnormal DNAme of miRNAs was involved in the development and progression of lung cancer. The data also suggested that targeting miRNA expression can be a therapeutic strategy to inhibit tumor progression and to improve patient survival.

#### Identification of methylation-driven lncRNAs in human lung cancer

To demonstrate the lncRNA mode of EpiMix, we used the same lung adenocarcinoma dataset^41^, and we aimed to identify differentially methylated lncRNA genes in tumors compared to normal tissues. Compared to protein-coding genes, lncRNAs are shorter, lower-expressed, less evolutionarily conserved, and expressed in a more tissue-specific manner^53^. To precisely quantify lncRNA expression from RNA-Seq, we used our previously developed pipeline^54^. With this pipeline, we combined the transcriptome annotations from GENCODE and NONCODE^55^. Raw sequencing reads were aligned to the combined transcriptome reference and quantified using the Kallisto-Sleuth algorithm^56,57^. Using this pipeline, we were able to detect the expression of 2,475 lncRNAs in both tumors and normal tissues. This number was three times higher compared to the lncRNAs detected by the traditional STAR-HTSeq pipeline. We then computationally deconvoluted bulk DNAme data and lncRNA expression data to cell-type-specific data (**Supplementary Fig. 3**). Since over 95% of the functional differential DNAme was found in epithelial cells (**Supplementary Fig. 4c-d**), we next focused our analysis on epithelial cells.

EpiMix identified 397 CpGs functionally associated with the expression of 132 lncRNAs in epithelial cells (**Fig.6a** and **Supplementary Table 8**). Of these CpGs, 146 (36.8%) CpGs associated with 69 genes were hypomethylated and 187 (47.1%) CpGs associated with 73 genes were hypermethylated. Seventy-two percent (72.0%) of the functional CpGs were located at the promoters, and this proportion was significantly higher than randomly selected CpGs (**Supplementary Fig.1e**, Fisher’s exact test, *P* < 0.0001). The differentially methylated regions were enriched with histone modifications marking for actively transcribed promoters and enhancers, including H3K27ac, H3K4me1 and H3K4me3 (**Supplementary Fig.1f**).

**Fig. 6.**
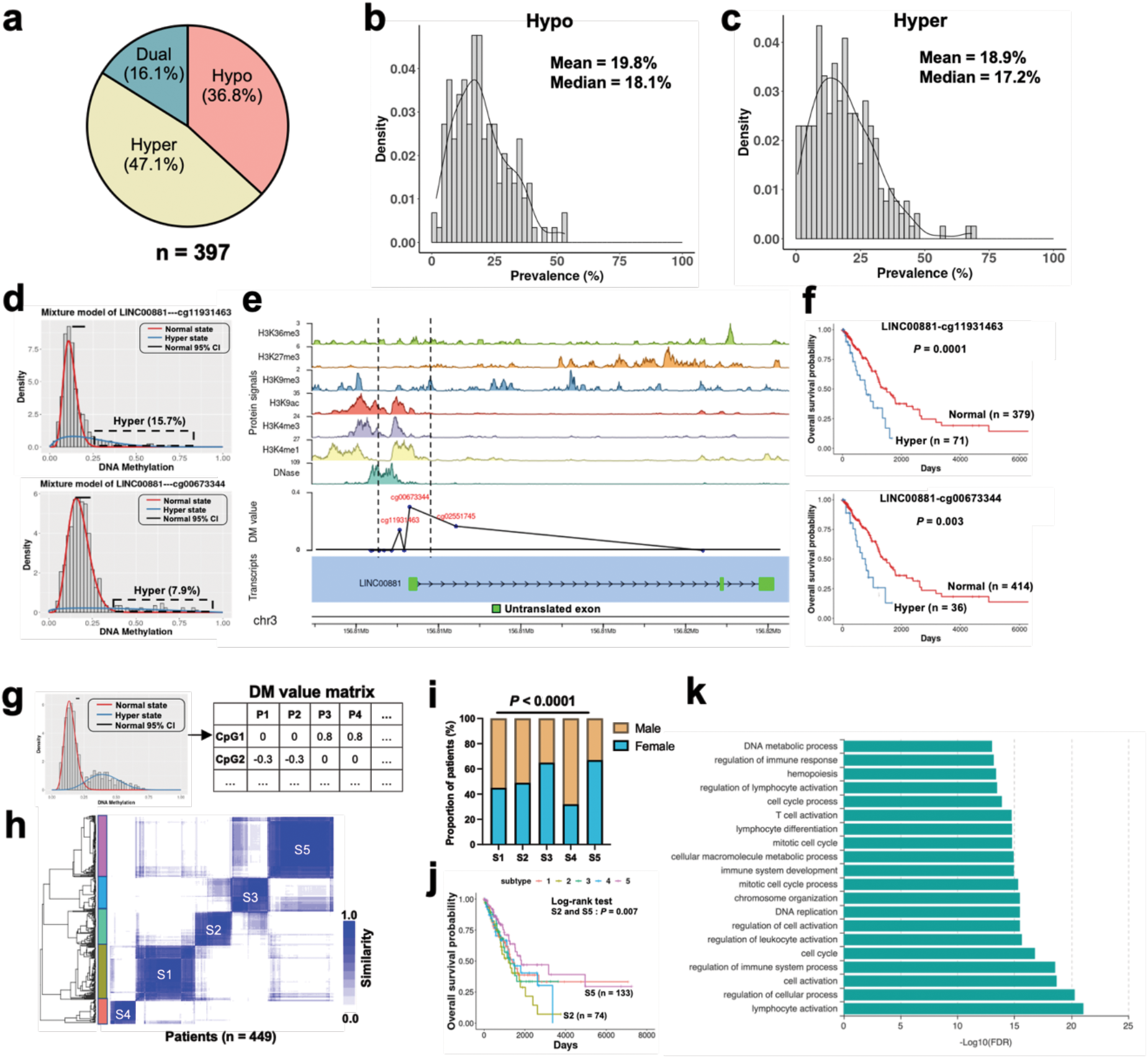
Identifications of differentially methylated lncRNA-coding genes in human lung cancers. **a**, Proportions of the hypo-, hyper- and dual methylated CpGs of lncRNA genes in epithelial cells from lung cancers compared to normal tissues. **b-c**, Density plot showing the prevalence distribution of the **(b)** hypo- and **(c)** hyper-methylated lncRNAs in the lung cancer cohort (n = 457). **d**, Mixture models of the *LINC00881* gene at two different CpG sites. Red indicates normal methylation and blue indicates hypermethylation. **e**, Integrative visualization of the transcript structure, DM values and chromatin state associated with the *LINC00881* gene. DM = 0: normal methylation; DM > 0: hypermethylation. **f**, Kaplan-Meier survival curves of patients in the normally methylated and the hypermethylated mixtures. Red indicates normal methylation and blue indicates hypermethylation. **g**, Schematic representation of the DM value matrix. The rows correspond to CpG sites, and the columns correspond to patients. DM values represent the mean differences in DNAme levels between patients in each mixture component identified in the experimental group compared to the control group. At each CpG site, patients in the same mixture component have the same DM values. DM < 0: hypomethylation, DM = 0: normal methylation, DM > 0: hypermethylation. **h**, Consensus matrix showing patient clusters based on the DM values of lncRNAs. **i**, Proportions of male and female patients in different patient clusters (n1 = 120, n2 = 74, n3 = 72, n4 = 50, n5 = 133). **j**, Kaplan-Meier survival curves of patients in different patient clusters. **k**, Top 20 enriched GO terms of the methylation-driven lncRNAs in lung cancer. DM: differential methylation.

The majority of differential methylation was identified in less 50% of the patients. The prevalence for hypomethylation ranged from 1.8% to 53.0%, with a mean value of 19.8% (**Fig.6b**). Similarly, the prevalence for hypermethylation ranged from 0.6% to 68.2%, with a mean value of 18.9% (**Fig.6c**). For instance, one of the hypermethylated lncRNAs was *LINC00881. LINC00881* was hypermethylated at CG11931463 in 15.7% of the patients and CG00673344 in 7.9% of the patients (**Fig.6d**). Both CpGs were located within the promoter (**Fig.6e**). Integrative analysis with clinical data showed that *LINC00881* hypermethylation was associated with significantly worse patient survival (**Figs.6f**, log-rank test, *P* < 0.001). These data demonstrated that many lncRNAs were differentially methylated in only a subset of the lung cancer patients. In addition, EpiMix was able to identify survival-associated lncRNAs that were differentially methylated in small patient subsets.

One of the major outputs from EpiMix is a differential methylation or “DM” value matrix, which reflects the homogeneous subpopulations of samples with a particular methylation state (**Fig.6g**). An application of the DM value matrix is to identify DNAme-associated subtypes, where patients are clustered into robust and homogenous groups based on their differential DNAme profiles. Using unsupervised consensus clustering, we discovered five DNAme subtypes (S1–S5) (**Fig.6h**). S5 contained a significantly higher proportion of females (89/133 = 66.9%) compared to S1 (54/120 = 45.0%), S2 (36/74 = 48.6%) and S4 (16/50 = 32.0%) (**Fig.6i**, Fisher’s exact test, *P* < 0.01). In addition, patients from S5 had significantly better survival than patients of S2 (**Fig.6j**, log-rank test, *P* = 0.007). We benchmarked the clustering results from using the DM value matrix versus using the raw DNAme data (beta values) of the differentially methylated CpGs. The patient subsets identified using raw DNAme data had low cluster consensus (**Supplementary Fig.5**), and no significant association was found between patient subsets and survival outcome. These results demonstrated that the DNAme subtypes discovered by EpiMix had prognostic values.

To investigate the biological functions of the differentially methylated lncRNAs, we utilized ncFANs, a functional annotation tool for lncRNAs^58^. We identified 4,552 protein-coding genes functionally associated with 76 lncRNAs. GO analysis showed that the protein-coding genes were primarily associated with DNA replication, cell cycle and regulation of cell activation (**Fig.6k** and **Supplementary Table 9**). These results indicated how differential methylation of lncRNAs were involved in the regulation of lung cancer development and progression.

## Discussion

In this study, we present EpiMix, a comprehensive analytic framework for population-level analysis of DNAme and gene expression. We packaged the EpiMix algorithms in R, enabling end-to-end DNAme analysis. To enhance the user experience, we also implemented a web-based application (https://epimix.stanford.edu) for interactive exploration and visualization of EpiMix’s results (**Fig.7**). EpiMix contains diverse functionalities, including automated data downloading, preprocessing, methylation modeling and functional analysis. The seamless connection of EpiMix to data from the TCGA program and the GEO database enables DNAme analysis on a broad range of diseases. Here, we showed that EpiMix identified novel methylation-driven pathways in food allergy and lung cancer. However, EpiMix is not limited to these disease areas and can be easily applied to any other diseases.

**Fig. 7.**
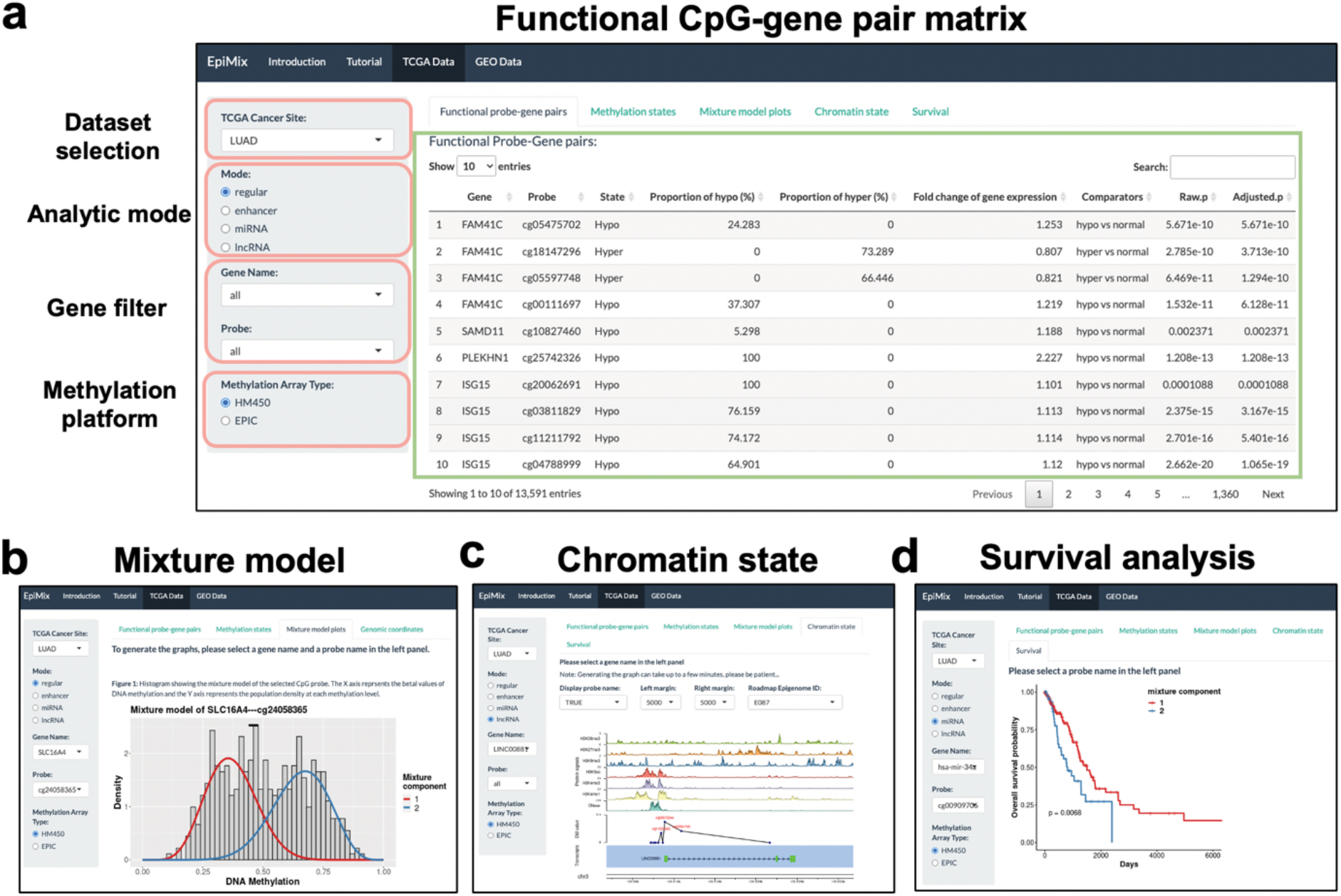
Screenshots of the EpiMix web application. **a**, Interactive data filters and visualization of functional CpG-gene pair matrix. **b**, Visualization of the mixture model of the SLC16A4 gene in lung cancer. **c**, Genome-browser style visualization of the lncRNA gene LINC00881 in lung cancer. **d**, Kaplan-Meier survival curves of patients with different methylation states of the miRNA gene miR-34a in lung cancer.

EpiMix uses a beta mixture model to decompose the DNAme profiles in a patient population. Using EpiMix, we can resolve the epigenetic subtypes within the patient population and pinpoint the individuals carrying differential DNAme profiles. In this study, we identified five DNAme subtypes in lung cancers using the DM values of lncRNAs. Patients of subtype 2 had worse survival than patients of subtype 5, indicating that the DNAme subtypes discovered by EpiMix had prognostic values. The biological interpretation of DNAme subtypes requires the integration of data from other modalities, such as genetic mutations, lifestyle history, and other etiological features.

In addition, EpiMix was able to detect abnormal DNAme that was present in only small subsets of a patient cohort. In our simulation study, EpiMix detected more differentially methylated CpGs compared to existing methods, when the differential methylation occurred in only a small patient subset. Using the real lung cancer dataset (n = 457), we identified miRNAs that were differentially methylated in only 1.1% of the patient population and lncRNAs differentially methylated in 0.6% of the patient population. We showed that over half of the miRNAs and lncRNAs were differentially methylated in only less than 20% of the patients. This unique feature of EpiMix to detect differential DNAme in small patient subsets enables us to identify novel epigenetic mechanisms underlying disease phenotypes. It can also be used to discover new epigenetic biomarkers and drug targets for improving personalized treatment.

Another feature of EpiMix is its ability to model DNAme at functionally diverse genomic elements. This includes *cis*-regulatory elements within or surrounding protein-coding genes, distal enhancers, and genes encoding miRNAs and lncRNAs. To model DNAme at distal enhancers, we selected the enhancers from the ENCODE and ROADMAP consortiums, in which enhancers of over a hundred human tissues and cell lines were identified using the chromatin-state discovery (ChromHMM)^59^. Since enhancers are cell-type specific, EpiMix allows the users to select enhancers of specific cell types or tissues. In this study, we selected the enhancers of human blood and T cells, leading to the discovery of 40,311 CpG of enhancers. In addition to enhancers, many other regulatory elements were identified from the ROADMAP studies^59^. These include active transcription start site proximal promoters, zinc finger protein genes, bivalent regulatory elements, polycomb-repressed regions and many others. By customizing the “chromatin state” parameter of EpiMix, users can target the DNAme analysis to any of these regulatory modules.

Despite the critical biological functions of non-coding RNAs, there are no existing tools that specifically analyze DNAme regulating their transcription. To analyze DNAme of miRNA genes, we utilized the miRNA annotation from miRBase, the largest and consistently updated knowledge base of miRNAs^60^. In addition, we selected CpGs at miRNA promoters by using a recent database that integrates the information of miRNA TSSs from 14 genome-wide studies across different human cell types and tissues^61^. This led to the discovery of 17,192 CpGs associated with 1,484 miRNAs in the HM450 array and 23,379 CpGs associated with 1,759 miRNAs in the EPIC array. With miRNA-Seq data provided, EpiMix can select differential DNAme that was associated with miRNA expression. Different from profiling protein-coding gene expression, measuring miRNA expression requires special library preparation strategies that capture small RNAs from total RNAs^62^. Users are preferentially needed to supply miRNA expression data obtained from proper library preparation strategies.

Similarly, custom methods are needed to accurately quantify lncRNA expression from RNA-Seq. We adopted the data processing pipeline developed from our previous study^54^. With this pipeline, we combined the transcriptome annotations from GENCODE and NONCODE. Raw sequencing reads were aligned to the combined transcriptome reference and quantified using the Kallisto-Sleuth algorithm^56,57^. Using this pipeline, we detected the expression of over 2,400 lncRNA genes. In this study, we have used our pipeline to generate lncRNA expression profiles for all the cancers in the TCGA database, and users can retrieve these data with EpiMix. Note, if users plan to use EpiMix on non-TCGA datasets, they are encouraged to use this pipeline to profile lncRNA expression.

Future work will aim to extend the use of EpiMix to whole-genome bisulfite sequencing and to further improve the scalability. Furthermore, the rapid development of single-cell technologies enables co-assay of DNAme and gene expression in thousands of cells. EpiMix can be used to identify differential DNAme that was present in only small subsets of a cell population. Therefore, a joint analysis of single cell methylome and transcriptome holds great promise for substantiating our goals, and the analytical framework presented here will be a valuable component for future research and applications.

## Methods

### Data downloading

The downloading module enables automated data downloading from the GEO database and TCGA project. Alternatively, users can supply custom datasets generated from their own studies. To retrieve data from GEO, we utilized the *getGEO* function from the GEOquery R package (version 2.62)^63^. In this study, we downloaded DNAme data and gene expression data using GEO accession number GSE114135. The DNAme data were beta values ranging from 0 to 1, representing the proportion of the methylated signal to the total signal. The gene expression data were TMM values. Other formats of gene expression data are also acceptable (e.g., RPKM, TPM, FPKM etc.). To retrieve data from TCGA, we used the Broad Institute Firehose tool (Firehose)^64^. We downloaded level three DNAme data and gene expression data. The downloaded data have been preprocessed for several steps, including removing problematic rows, removing redundant columns, reordering the columns and sorting the data by gene name. With the Regular mode, we used log-transformed RSEM values. With the miRNA mode, we used the pri-miRNA expression data with log-transformed RPKM values.

### Preprocessing

The majority of datasets obtained from the TCGA and GEO databases have already been preprocessed for a few steps. EpiMix’s contribution to preprocessing includes missing value imputation, removal of single-nucleotide polymorphism (SNP) probe and batch effect correction. Users can also select to remove CpGs on sex chromosomes. We then removed CpGs and samples with more than 20% missing values, and imputed missing values on the remaining dataset using the k-nearest neighbor (KNN) algorithm with K = 15.

Data from large patient cohorts were typically collected in technical batches. Systematic variances between technical batches may affect downstream data analysis and interpretation. To correct batch effects, we implemented two alternative approaches: (1) an anchor-based data integration approach adapted from the Seurat package (version 4.0.1)^65^ and (2) an empirical Bayes regression approach, Combat^66^. The anchor-based approach uses canonical correlation analysis and mutual nearest neighbors to identify shared subpopulations (termed “anchors”) across different datasets and then uses a non-linear transformation to integrate the data. To identify the anchors, we used the “vst” method to select the top 10% variable features. Effective batch effect removal was confirmed using the PCA-based ANOVA analysis. Alternatively, the batch effect can be corrected with the Combat algorithm^58^. We found that the anchor-based approach was more time efficient compared to the Combat. When tested on the lung cancer dataset, the former approach completed the batch correction within 2 hours, whereas the Combat consumed more than 48 hours.

### CpG annotation and filtering

#### Regular mode

The Regular mode aims to model DNAme at cis-regulatory elements within or immediately surrounding protein-coding genes. We paired each CpG site to the nearest genes based on the hg38 manifest generated from Zhou et al.^67^. Unique CpG-gene pairs were identified, where a CpG was either within the gene body or at the immediately surrounding area. Users can restrict the analysis to the promoters, defined as 2 kb upstream and 500 bp downstream (-2000bp ∼ +500bp) of the transcription start sites (TSSs). TSS information was retrieved from Ensembl using the *biomaRt* R package (version 2.50.1)^68^.

#### Enhancer mode

The Enhancer mode aims to model DNAme specifically at distal enhancers. Therefore, we selected the distal CpGs that were at least 2 kb away from any known TSSs. Users can customize this distance based on their needs. To select the CpGs within enhancers, we used the enhancer database established from the ENCODE and ROADMAP consortiums, in which enhancers of over a hundred human tissues and cell lines were identified using the chromatin-state discovery (ChromHMM)^59^. We looked for the DNA elements associated with the chromatin states of active enhancers (“EnhA1” and “EnhA2”) and genic enhancers (“EnhG1” and “EnhG2”). Since enhancers are cell-type specific, EpiMix allows users to select enhancers of specific cell types or tissue groups. In this study, we selected the enhancers of human blood and T cells, leading to the discovery of 40,311 CpGs of enhancers. For each CpG, we retrieved 20 nearby genes as candidate genes targets. This gene number was determined by the previous studies showing that many of the enhancers can regulate a gene within a 10-gene distance^29,69,70^. Genes that are positively regulated by the enhancers should have a negative relationship between DNAme and gene expression^36,71,72^. Therefore, we performed a one-tailed Wilcoxon rank-sum test on each enhancer-gene pair to select the enhancers whose methylation states were inversely associated with the gene expression. The raw *P* value from the Wilcoxon rank-sum test was adjusted using a permutation approach^73^, where an empirical *P* value was determined by ranking the raw *P* value in a set of permutation *P* values from testing the expression of a set of randomly selected 1,000 genes (**Supplementary Fig.2**).

#### miRNA mode

MicroRNAs are commonly classified into “intergenic” or “intronic” based on their genomic locations. Intergenic miRNAs are found at previously unannotated human genome and are transcribed from their own unique promoters as independent entities. In contrast, intronic miRNAs are believed to share promoters with their host genes and co-transcribed from respective hosts. Recent evidence shows that some intronic miRNAs can also be transcribed independently from their host genes, suggesting they have their own independent promoters^74^. To select CpGs associated with miRNAs, we used a combined strategy. First, we obtained the most recent annotation of miRNAs from miRBase (version 22.1)^60^. For each miRNA gene, we selected CpGs that were located within 5 kb upstream and 5 kb downstream. Second, we selected CpGs at miRNA promoters by using a recent database that integrates miRNA TSS information from 14 genome-wide studies across different human cell types and tissues^61^. We included CpGs located with miRNA promoters defined as 2000 bp upstream and 1000 bp downstream of the TSSs. This combined feature selection strategy resulted in the discovery of 17,192 CpGs associated with 1,484 miRNAs in the HM450 array and 23,379 CpGs associated with 1,759 miRNAs in the EPIC array.

#### lncRNA mode

The mechanisms for transcriptional regulation of lncRNAs are similar to protein-coding genes. We first selected lncRNA-coding genes using the GENCODE annotation (Version 36). We then selected CpGs associated with each lncRNA based on the hg38 manifest generated from Zhou et al.^67^. Unique CpG-gene pairs were identified, where a CpG was either located within the gene body or at the immediately surrounding area. This resulted in the discovery of 98,320 CpGs associated with 11,280 lncRNAs in the HM450 array and 184,816 CpGs associated with 15,392 lncRNAs in the EPIC array. Alternatively, users can select to focus the analysis at lncRNA promoters, defined as 2 kb upstream and 500 bp downstream (-2000bp ∼ +500bp) of the TSSs. The TSS information was retrieved from Ensembl using the *biomaRt* R package (version 2.50.1)^68^.

### CpG site clustering and smoothing (optional features)

#### Clustering

Modeling the DNAme at all individual CpG sites can be computationally expensive. In addition, it can also lead to overfitting of DNAme data in identifications of patient subsets. Since the DNAme at adjacent CpGs are strongly correlated, we implemented an optional feature that allows users to group the correlated CpGs into CpG clusters. First, we used the average linkage hierarchical clustering algorithm to cluster CpGs of a single gene into clusters. Then we cut off the hierarchical tree at a Pearson correlation threshold of 0.4 to define CpG clusters and single CpG sites when they do not correlate with other sites. For each CpG site cluster, we used the mean levels of DNAme of the CpGs to represent the cluster DNAme, resulting in potentially multiple CpG site clusters representing a single gene. The DNAme modeling can then be performed at each separate CpG site or CpG site cluster.

#### Smoothing

Smoothing is another technique frequently used in removing noise and increasing statistical power in analyzing whole-genome bisulfite sequencing data^6^. This technique estimates localized DNAme levels using data of adjacent CpGs at a user-specified genomic window. EpiMix allows users to smooth the DNAme data using local likelihood smoothing^75^. Since the number of CpGs is lower in array-based data than in bisulfite sequencing data, using smoothing on array-based data should be taken with cautions.

### Methylation modeling

After preprocessing, the methylation data are beta values bounded between 0 and 1, representing the proportion of the methylated signal to the total signal. When the study population is large, the beta values can be assumed to come from multiple underlying probability distributions, in our case, beta distributions. To model the DNAme, we fit a beta mixture model to the methylation values at each CpG site (or CpG site cluster). Let *y*_*i*_ denote the beta value from subject *i* at a CpG site, where *i* ∈ {1,…, *n*}, and *n* represents the total number of subjects. Let *k* denote the class membership of subject *i*, where *k* ∈ {1,…, *K*}, and *K* represents the total number of components in the mixture.

Assume subject *i* belongs to component *k* with probability *η*_*k*_, we will have 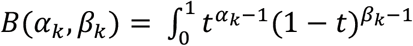. Subsequently, the likelihood contribution from subject *i* is:

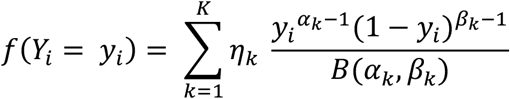

where 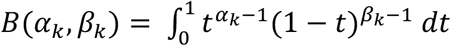 is the beta function. Since the population contains *n* subjects, the log-likelihood for the complete dataset is

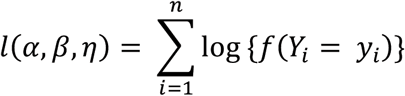

The goal of our modeling is to estimate the *α, β, η* parameters of each component that best fit the methylation values. Let *θ* = {*α*_1,_ *β*_1,_ *η*_1_…, *α*_*k*,_ *β*_*k*_, *η*_*k*_} be a vector of parameters that define the shape of each component in the mixture. We used the expectation–maximization (EM) algorithm^76^ to iteratively maximize the log-likelihood and update the conditional probability that *y*_*i*_ comes from the *k th* component.

To determine the best number of components *K*, we used The Bayesian Information Criterion (BIC) for model selection and to avoid overfitting:

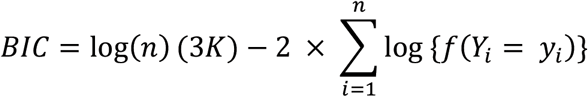

This process involves iteratively adding a new mixture component if the BIC improves. Each mixture component represents a subset of samples for whom a particular DNAme state is observed.

### Identifications of differentially methylated CpGs

If data of a control group are provided, we can determine whether a CpG site (or CpG site cluster) was hypo- or hyper-methylated by comparing its methylation levels in the experimental group to its counterpart in the control group. We first performed beta mixture modeling on each CpG site (or CpG site cluster) to identify the mixture components using data from the experimental group, and the methylation levels of each of the mixture components were compared to the mean methylation levels of the control group. This methodology is based on the assumption that the DNAme profile is heterogenous across different subjects in the experimental (i.e., disease) group but is homogenous in the control group. For instance, the DNAme profile is expected to be different across cancer patients due to the difference in subtypes or driver mutations, but in normal tissues the DNAme should be relatively homogenous. In addition, the number of subjects in the experimental group is typically higher than the control group (e.g., TCGA projects). To determine the significant difference between the experimental and the control group, we used a Wilcoxon rank-sum to calculate the *P*-value, and multiple comparison was corrected with the false discovery rate (FDR). The Q-value threshold was set to 0.05. In addition, we required a minimum difference of 0.10 based on the platform sensitivity reported previously^77^.

### Identifications of differential DNAme that was associated with transcription

If sample-matched gene expression data are provided, we can select the CpGs whose methylation states were significantly associated with gene expression. In this study, we focused on the identification of DNAme that represses gene expression. However, users have the option to identify DNAme that is positively correlated with gene expression. For each CpG-gene pair, we used a one-tailed Wilcoxon rank-sum test to compare the mean levels of gene expression in patients showing an abnormal methylation state (hypo- or hyper-methylation state) to those with a normal methylation state. If a CpG was hypomethylated, we examined that the hypomethylated patients have higher gene expression levels compared to the normally methylated patients. Vice versa, if a CpG was hypermethylated, we tested that the hypermethylated patients have lower gene expression levels compared to the normally methylated patients. If a CpG was dual methylated (i.e., some samples were hypomethylated, while some others were hypermethylated), we tested that the hypomethylated patients have higher gene expression levels compared to the hypermethylated patients. Since a gene is typically paired with multiple CpGs, we adjusted the *P*-value using FDR to correct multiple comparisons. To select functionally significant CpG-gene pairs, we set the maximum threshold of the adjusted *P*-value to 0.01.

### Simulation study

The goal of the simulation studies was to assess the sensitivity of EpiMix to detect differential DNAme present in only specific subsets of a population. The studies were performed by creating synthetic CpG sites and synthetic populations. First, we filtered CpGs showing statistically similar DNAme levels that fit a unimodal beta distribution from the activation group and from the quiescent group (n = 103 samples per group). We then randomly sampled a subset of CpGs (n = 300) from the quiescent group as the baselines. The average DNAme levels (beta values) of the CpGs in the baseline group ranged from 0.1 to 0.9, with a mean DNAme level of 0.6. Second, since the magnitude of changes in DNAme levels can be a critical factor affecting sensitivity, we created synthetic CpGs. For each CpG of the baseline group, we paired it with a subset of CpGs from the activation group, such that the differences in the mean beta values (*Δbeta*) between the the activation group and the baseline group ranged from 0.1 to 0.7, where *Δbeta* ∈ {0.10, 0.15, 0.20, 0.25, 0.30, 0.40, 0.50, 0.60, 0.70}. This resulted in a total of 2,700 synthetic CpGs. Third, since our goal was to detect differential DNAme that was present in only a subset of the population, we created synthetic populations. For each synthetic CpG, we controlled the number of samples from the activation group to be combined with the baseline group, such that the final proportion (*P*) of samples from the activation group in the combined datasets ranged from 0.01 to 0.50, where *P* ∈ {0.01, 0.02, 0.05, 0.08, 0.10, 0.15, 0.20, 0.25, 0.30, 0.35, 0.40, 0.45, 0.50}. Finally, we ran the EpiMix algorithm on each synthetic CpG and assessed whether it could pick up the differentially methylated signals in the synthetic populations.

### Benchmark with existing methods

We benchmarked the performance of EpiMix with other existing methods, including Minfi^10^, iEVORA^27^ and RnBeads^12,28^.

Minfi includes a differential methylation step based on an F-test. We first transformed beta values to M values, and the differential methylation analysis was performed with the *dmpFinder* function. We set the significant *P*-value and *Q*-value thresholds to 0.05.

iEVORA is a two-step algorithm that selects differentially variable and differentially methylated CpGs. The first step is to identify differentially variable CpGs using a Bartlett’s test. The Bartlett’s test assesses the equity of variances between the experimental and the control group. If in the experimental group, there are samples showing large differences (outliers) in DNAme versus other samples, the Bartlett’s test can detect such abnormality. The second step is to select the differentially variable CpGs that were also differentially methylated. The differential methylation analysis is performed by comparing the mean levels of DNAme of all the samples in the experimental group to the control group. We used the default parameters of the functions, with a *Q*-value (FDR) threshold of 0.001 for testing differential variability and *P*-value threshold of 0.05 for testing differential methylation means. In our stimulation studies, we found that iEVORA was able to identify differentially variable CpGs even when the abnormal methylation was present in only a small subset of the experimental group. However, since the algorithm does not identify which subjects were abnormally methylated, and in the differential methylation step, it still compares the mean levels of DNAme of the entire experimental group to the control group, the differential methylation test could not generate statistically significant results.

RnBeads uses hierarchical linear models as implemented in the limma package to identify differential methylated CpGs. We set the differential methylation *P*-value threshold to 0.05.

### Imputation of cell-type-specific DNAme and gene expression data

DNAme and gene expression are known to be cell-type specific. When the DNAme were measured at the tissue (“bulk”) level, the differential DNAme profiles between patient subjects may result from the differences in tissue compositions. From a clinical perspective, tissue composition is meaningful in classifications of tumor subtypes and prediction of treatment response. However, from a biological perspective, users may be interested in identifying the differential DNAme present in specific cell types. EpiMix focuses on the identification of differential DNAme across patient individuals. To resolve the confounding effect from tissue heterogeneity, we used previously validated algorithms to infer cell-type proportions and cell-type specific methylomes and transcriptomes (**Supplementary Fig.3**). First, we used CIBERSORTx^45^, a reference-based computational algorithm, to estimate cell-type proportions from bulk gene expression data in each tumor and normal tissue, and deconvolute bulk gene expression data into cell-type specific signals. This method leveraged the established signature gene expression matrices for experimentally purified cells from normal tissues and lung cancers^45^. Second, we used Tensor Composition Analysis (TCA)^44^ to deconvolute bulk DNAme data into cell-type-specific data based on the estimated cell-type proportions in each tissue. The output from TCA was the methylome of each cell type in each individual. In addition to these methods, users can leverage other existing tools to adjust the effects from tissue compositions before inputting the data to EpiMix^78–83^.

### Genomic distribution of the differentially methylated CpGs

Genomic coordinates of the TSSs of the methylation-driven genes were retrieved from Ensembl using the *biomaRt* R package (version 2.50)^68^. Exons and Introns of the protein-coding genes were retrieved from the TxDb object (*TxDb*.*Hsapiens*.*UCSC*.*hg38*.*knownGene*) (version 3.14)^84^. The GenomicRanges R package (version 1.46)^85^ was used to identify the differentially methylated CpGs located within promoters, exons and introns.

### Motif enrichment analysis

TF binding motifs were retrieved from HOCOMOCO, a comprehensive database for TF binding sites^86^. HOMER (Hypergeometric Optimization of Motif EnRichment) was used to find motif occurrences in a ±250bp region around each differentially methylated regions (DMRs). We then combined all the DMRs to identify enriched motifs. Enrichments were quantified using Fisher’s exact test and multiple comparisons were adjusted with the Benjamini-Hochberg procedure. To calculate the enrichment Odds Ratio, we used all the distal CpGs as the background probes and the functional CpGs of enhancers as the target probes. We set the significant *P* value cutoff to 0.05 and the smallest lower boundary of 95% confidence interval for Odds Ratio to 1.1. The enrichment analysis was performed using the *get*.*enriched*.*motif* function from the *ELMER* library (version 3.14) in R^11^.

### Enrichment analysis of chromatin modifications

Enrichment analysis of histone modifications at the DMRs was performed using the Genomic Hyperbrowser GSUITE of tools^87^. A suite of tracks representing different chromatin features for human naïve T cells (Epigenome ID: E038) and lung (Epigenome ID: E096) were retrieved from the ENCODE and ROADMAP consortiums^59^. To determine which tracks in the suite exhibit the strongest similarity by co-occurrence to the DMRs, the Forbes coefficient was used to obtain rankings of tracks, and Monte Carlo simulations were used to define a statistical assessment of the robustness of the rankings using randomization of genomic regions covered by the entire HM450 or EPIC array, and compute test statistics.

### Functional enrichment analysis

#### Protein-coding genes

EpiMix provides an user interface to the *enrichGO* and *enrichKEGG* functions of the *clusterProfiler* R package (version 4.2.1)^88^. This enables gene set analysis of the methylation-driven genes using the gene ontology (GO) and KEGG datasets. Over-represented biological pathways in the methylation-driven genes were identified using the hypergeometric testing^88^. Enrichment results can be retrieved in a tabular format or visualized in several different ways. To perform the GO analysis, we set the significant *P* value to 0.05 and Q value to 0.20. Highly similar GO terms were removed with a cutoff *P* value of 0.60 to retain the most representative terms.

#### miRNAs

To obtain the target genes of the differentially methylated miRNAs, we queried miRTarBase with the *miRnetR* package^89^. Of the 144 differentially methylated miRNAs in lung cancer, we identified 7,088 target protein-coding genes of 26 miRNAs. We simplified this network by selecting the genes that were targeted by at least five miRNAs. KEGG pathway analysis was then performed on the miRNA target genes with hypergeometric testing.

#### lncRNAs

To carry out functional annotation and pathway analysis of the differentially methylated lncRNAs, we used the ncFANs V2.0 server (http://ncfans.gene.ac/)^58^. The genes in the significant CpG-gene pair matrix generated from EpiMix can be directly used as an input to ncFANs. NcFANs assigns the functions of protein-coding genes to lncRNAs based on pre-built co-expression networks in various normal tissues and cancers. We used the co-expression network built in the lung adenocarcinoma dataset from TCGA, and we set the correlation coefficient between lncRNAs and proteins-genes to 0.4 and the cutoff of the topological overlap measure similarity to 0.01.

### Biomarker identification and survival analysis

Patient clinical data were retrieved from TCGA using the Firehose tool^64^. Alternatively, users can provide EpiMix with survival data if using their own datasets. We selected the CpGs with at least two methylation states. For each CpG, we fit a Cox proportional hazards regression model to assess the effect of methylation states on patient survival time. The log-rank test was used to compare the survival curve and to calculate the significant *P*-value. *P* < 0.05 was considered as significant. The Kaplan-Meier survival plots were generated with the *survminer* R package (version 0.4.9).

### Genome browser-style visualization

EpiMix enables genome browser-style visualization of the genomic coordinates and chromatin states of the differentially methylated genes and regions. We implemented two different forms of visualization. The gene-centric form shows the DM values of all the CpGs associated with a specific gene (e.g., **Fig.3f)**. The CpG-centric form shows a differentially methylated CpG and its upstream and downstream genes (e.g., **Fig.4e)**. Users can specify the number of nearby genes to display. Genes whose expression levels were significantly associated with the DNAme levels of the CpG are shown in red.

DNase I sensitivity and histone modification levels were retrieved from the ENCODE and ROADMAP consortiums^59^. By providing the Epigenome ID, users can retrieve data corresponding to the investigated tissue or cell type. In this study, we extracted the chromatin features for human naïve T cells (Epigenome ID: E038) and fetal lung (Epigenome ID: E088). The genomic coordinates (X-axis) were established on the hg19 genome built, and the enrichment signal (Y-axis) represents negative log10 of the Poisson *P*-values. Human transcript annotation was retrieved from the TxDb object (*TxDb*.*Hsapiens*.*UCSC*.*hg19*.*knownGene*) (version 3.2.2)^90^. The genomic coordinates of the adjacent genes of the differentially methylated CpGs were retrieved from Ensembl using the *biomaRt* R package (version 2.50.1)^68^. The visualization was implemented with the *karyoploteR* package (version 1.20.0)^91^.

### Identifications of DNAme subtypes

DNAme subtypes can be discovered by applying consensus clustering to the DM-value matrix, where patients were clustered into robust and homogenous groups (putative subtypes) based on their abnormal methylation profiles. Consensus clustering was performed with the ConsensusClusterPlus R package (version 1.58.0)^92^. We used 1,000 rounds of k-means clustering and a maximum of K = 10 clusters. Selection of the best number of clusters was based on the visual inspection of ConsensusClusterPlus output plots.

## Code availability

EpiMix is available as an R package on Bioconductor (https://bioconductor.org/packages/devel/bioc/html/EpiMix.html). In addition, we also developed a web application (https://epimix.stanford.edu) for users to interactively visualize and explore the results from EpiMix.

## Acknowledgements

We thank Dr. Sandra Steyaert, Alexander Henry Thieme (Department of Radiation Oncology, Charité - Universitätsmedizin Berlin), and Gautam Machiraju for helpful comments and suggestions.

## Funding

Research reported here was further supported by the National Cancer Institute (NCI) under awards: R01 CA260271, U01 CA217851 and U01 CA199241. The content is solely the responsibility of the authors and does not necessarily represent the official views of the National Institutes of Health.

## Author contributions

Y.Z: study design, implementation, data analysis and interpretation of results, draft manuscript preparation J.J: implementation and data analysis K.B: study conception and design O.G: study conception and design, resources, supervision

## Competing interests

The authors declare no competing interests.

## Notes

### Competing Interest Statement

The authors have declared no competing interest.

