## Supplementary Figures for "EpiMix: an integrative tool for epigenomic subtyping using DNA methylation"

### Supplementary Fig.1

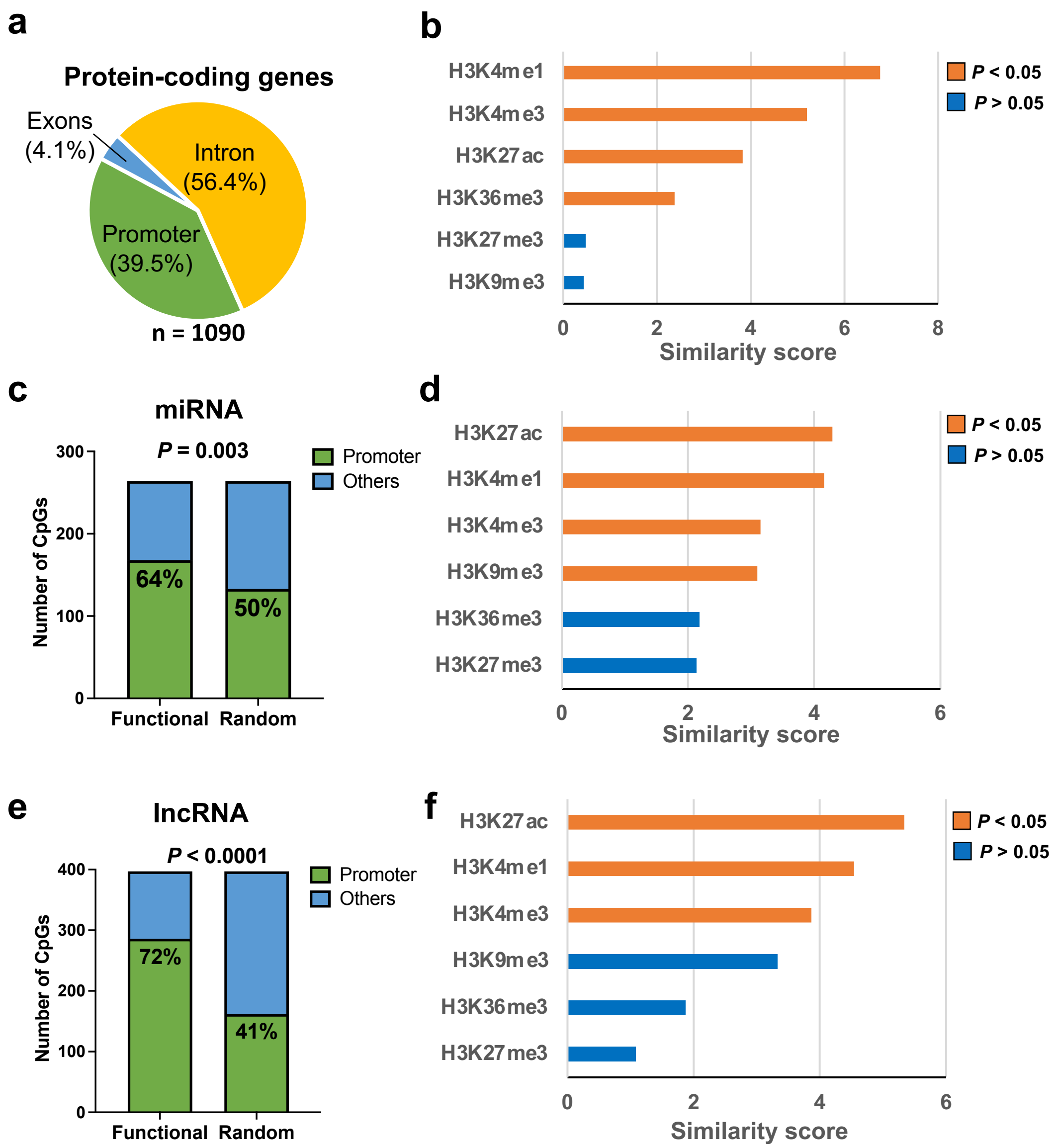


**Supplementary Fig.1: a,** Proportions of the functional CpGs at promoters (-2000 ~ +500bp flanking TSSs), exons and introns of protein-coding genes in antigen-activated T cells. **b,** Similarity (Forbes coefficient) between the differentially methylated regions (DMRs) and regions marked by specific histone marks. Similarity score was calculated by ratio of observed/expected overlap between the DMRs and the histone modification-enriched regions. Significant similarity ($P < 0.05$) was shown in orange. **c,** Proportions of the functional CpGs (n = 264) at the promoters and non-promoter regions of miRNAs. To investigate whether the functional CpGs were disproportionately located at the promoters, we randomly selected an equal number (n = 264) of CpGs from all the miRNA-associated CpGs and assessed how likely the random CpGs were located at the promoters. This experiment was repeated for 1,000 times, and the mean distribution was used. Fisher's exact test was used to compare the distributions of the functional CpGs and the randomly selected CpGs. **d,** Similarity (Forbes coefficient) between the DMRs of miRNAs and regions marked by specific histone marks. Similarity score was calculated by ratio of observed/expected overlap between the histone mark-enriched regions and the DMRs. Significant enrichment ($P < 0.05$) was shown in orange. **e,** Proportions of the functional CpGs (n = 397) at promoters and non-promoter regions of lncRNAs. To investigate whether the functional CpGs were disproportionately located at the promoters, we randomly selected an equal number (n = 397) of CpGs from all the lncRNA-associated CpGs and assessed how likely the randomly selected CpGs were located at the promoters. This experiment was repeated for 1,000 times, and the mean distribution was used. Fisher's exact test was used to compare the distributions of the functional CpGs and the randomly selected CpGs. **f,** Similarity (Forbes coefficient) between the DMRs of lncRNAs and regions marked by specific histone marks. Similarity score was calculated by ratio of observed/expected overlap between the histone mark-enriched regions and the DMRs. Significant enrichment ($P < 0.05$) was shown in orange.

### Supplementary Fig.2

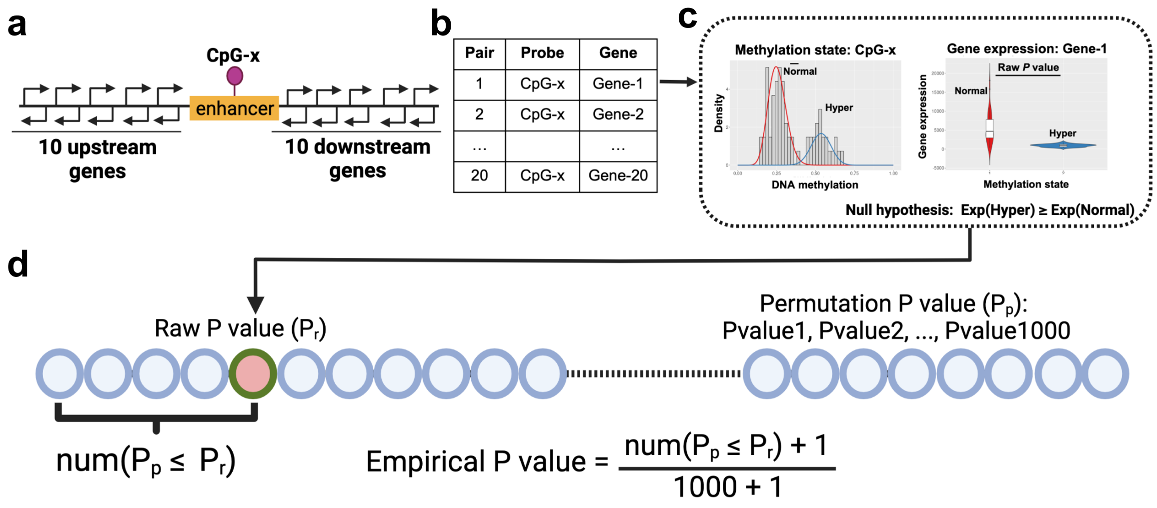


**Supplementary Fig.2: Strategy to identify the target genes of enhancers.** Enhancers of human blood and T cells were retrieved from the enhancer database established from the ENCODE and ROADMAP consortiums (see **Methods**). For each enhancer, we assigned 20 adjacent genes (i.e., 10 genes upstream and 10 genes downstream) as their candidate gene targets (**panel a**), which resulted in 20 unique enhancer-gene pairs (**panel b**). Genes that are positively regulated by the enhancer should have a negative relationship between the DNAm and gene expression. Therefore, we used a one-tailed Wilcoxon rank-sum test to examine whether the methylation state of an enhancer was inversely associated with the expression levels of each of its candidate gene target (**panel c**). This process generated a raw *P* value for each of the enhancer-gene pair. To adjust the *P* value, we assigned each enhancer with a set of 1,000 randomly selected genes, where we required the random genes to be located on randomly different chromosomes than the tested enhancer. We then performed the same Wilcoxon rank-sum test to examine whether the methylation states of the tested enhancer was inversely associated with the expression of each of the randomly selected genes. This generated 1,000 permutation *P* values (**panel d**). We then ranked the raw *P* value in the permutation *P* values to determine an empirical *P* value. We required the empirical *P* value to be smaller than 0.05 in order to select the target genes of an enhancer.

### Supplementary Fig.3

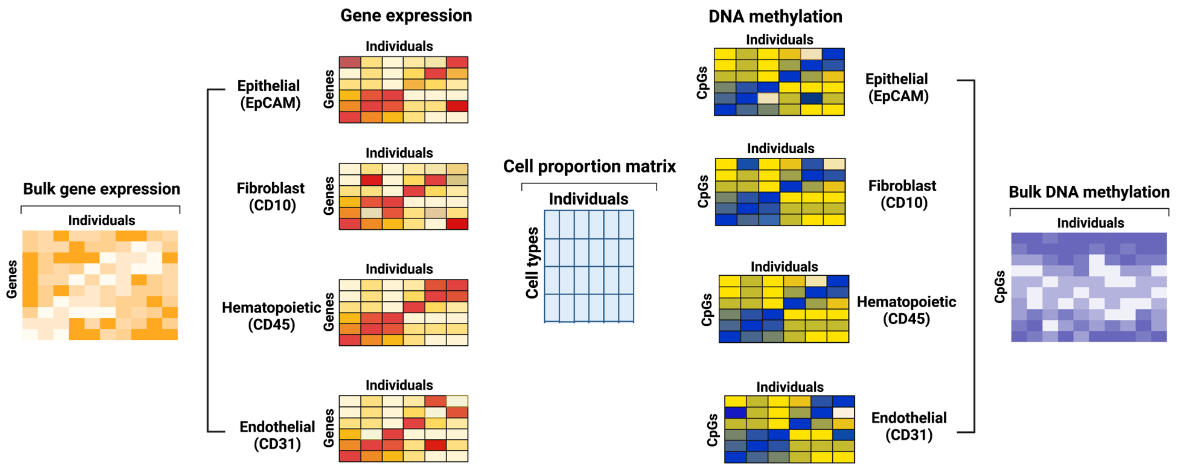


**Supplementary Fig.3: Strategy to deconvolute bulk DNAme data and gene expression data to cell-type-specific signals**. First, we used CIBERSORTx, a reference-based computational algorithm, to estimate cell-type proportions in each individual from the bulk gene expression data, and deconvolute bulk gene expression data into cell-type-specific signals. Second, we used Tensor Composition Analysis (TCA) to deconvolute bulk DNAme data into cell-type-specific data, based on the estimated proportion of each cell type. TCA generates the methylomes of each cell type in each individual.

### Supplementary Fig.4

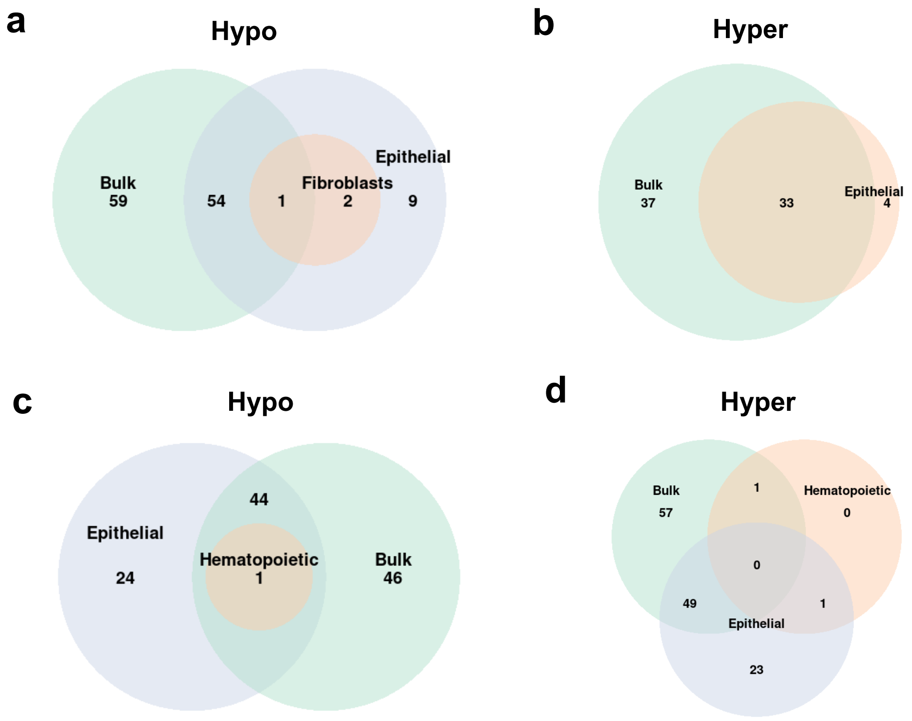


**Supplementary Fig.4: Venn diagrams showing the numbers of hypo- and hypermethylated genes identified using data from bulk tissues versus deconvoluted data of specific cell types.** DNA methylation data and gene expression data collected at tissue level ("bulk") were computationally deconvoluted into epithelial/cancer cells (EpCAM+), fibroblasts (CD10+), hematopoietic cells (CD45+) and endothelial cells (CD31+). **a,** numbers of hypomethylated miRNA genes identified in bulk tissues, epithelial cells and fibroblasts. **b,** numbers of hypermethylated miRNA genes identified in bulk tissues and epithelial cells. **c,** numbers of hypomethylated lncRNA genes identified in bulk tissues, epithelial cells and hematopoietic cells. **d,** numbers of hypermethylated lncRNA genes identified in bulk tissues, epithelial cells and hematopoietic cells.

### Supplementary Fig.5

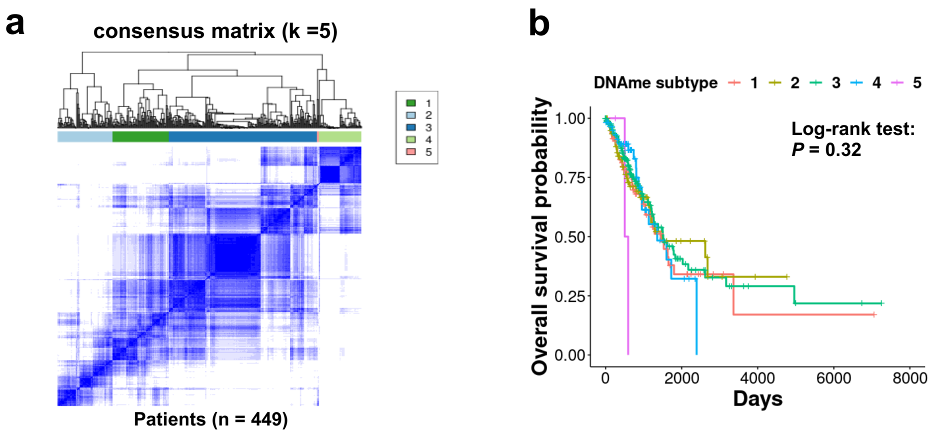


**Supplementary Fig.5: Survival analysis of patient subsets identified by consensus clustering of raw beta values. (a)** Consensus matrix showing patient clusters classified by raw beta values of the differentially methylated CpGs (n = 397). **(b)** Kaplan-Meier survival curves of patients of different DNAme subgroups (n1 = 84, n2 = 81, n3 = 221, n4 = 60, n5 = 3).
